# Agonist-dependent and -independent kappa opioid receptor phosphorylation showed distinct phosphorylation patterns and resulted in different cellular outcomes

**DOI:** 10.1101/137570

**Authors:** Yi-Ting Chiu, Chongguang Chen, Stefan Schulz, Lee-Yuan Liu-Chen

## Abstract

We reported previously that the selective agonist U50,488H promoted phosphorylation of the mouse kappa opioid receptor (KOPR) at residues S356, T357, T363 and S369. Here, we found that agonist (U50,488H)-dependent KOPR phosphorylation at all the residues were mediated by Gi/o_α_ proteins and multiple protein kinases [GRKs2, 3, 5 and 6 and protein kinase C (PKC)]. In addition, PKC activation by phorbol ester induced agonist-independent KOPR phosphorylation. Compared with U50,488H, PKC activation promoted much higher S356/T357 phosphorylation, much lower T363 phosphorylation and similar levels of S369 phosphorylation. Following U50,488H, GRKs, but not PKC, were involved in agonist-induced KOPR internalization. In contrast, PKC activation caused a lower level of agonist-independent KOPR internalization, compared to U50,488H. U50,488H-induced activation of extracellular signal regulated kinase 1/2 (ERK1/2) was G protein-, but not β-arrestin-, dependent. After U50,488H, GRK-mediated, but not PKC-mediated, KOPR phosphorylation followed by β-arrestin recruitment desensitized U50,488H-induced ERK1/2 response. Therefore, agonist-dependent (GRK- and PKC-mediated) and agonist-independent (PKC-promoted) KOPR phosphorylations show distinct phosphorylation patterns, leading to diverse cellular outcomes.

Abbreviations

CHO cells: Chinese hamster ovary cells
CHL: Chelerythrine chloride
DDM: Dodecyl-β-Dmaltoside
ERK1/2: extracellular signal regulated kinase 1/2
ECL: enhanced chemiluminescence
GRK: G protein-coupled receptor kinase
GPCRs: G protein-coupled receptors
HEK293 cells: human embryonic kidney 293 cells
HRP: Horseradish peroxidase
KOPR: kappa opioid receptor
MEM: Minimal essential medium
N2A cells: neuro2A mouse neuroblastoma cells
Ni-NTA: Nickel-nitrilotriacetate
NTF: no transfection
PKC: protein kinase C
PMA: phorbol-12-myristate-13-acetate
PTX: pertussis toxin

## Introduction

The kappa opioid receptor (KOPR) is one of the three opioid receptors and belongs to the rhodopsin subfamily of G protein-coupled receptors (GPCRs). Activation of the KOPR produces many effects, including analgesia, antipruritic effects, water diuresis, hypothermia, sedation, and dysphoria (Ansonoff et al., 2006; Cowan et al., 2015; Lemos and Chavkin, 2011; Simonin et al., 1998). At the cellular level, activation of KOPR stimulates Gi/o protein-mediated signaling and KOPR phosphorylation. Phosphorylated receptor recruits β-arrestin, leading to desensitization, internalization and β-arrestin-dependent signaling [reviewed in (Al Hasani and Bruchas, 2011; Law, 2011; Liu-Chen, 2004)]. Gi/o protein-dependent responses include inhibition of adenylyl cyclases and Ca^2+^ inward current, and activation of K^+^ channels and ERK1/2 (early phase) (Al Hasani and Bruchas, 2011; Law, 2011). β-arrestin-dependent signaling includes phosphorylation of ERK1/2 (late phase) and p38 MAP kinase (Al Hasani and Bruchas, 2011; Law, 2011).

Previous studies using different strategies have shown that agonists cause KOPR phosphorylation. We have demonstrated that U50,488H induces phosphorylation of the human KOPR expressed in Chinese hamster ovary cells (CHO cells) using metabolic labeling with ^32^P-orthophosphate (Li et al., 2002). Chavkin and colleagues reported that U50,488H induced rat KOPR (rKOPR) phosphorylation at S369 in human embryonic kidney 293 cells (HEK293 cells) by immunoblotting (McLaughlin et al., 2003). Recently, by liquid chromatography-tandem mass spectrometry (LC-MS/MS) analysis, we determined the phosphorylation sites of the mouse KOPR (mKOPR) following U50,488H treatment to be S356, T357, T363 and S369 in the C-terminal domain (Chen et al., 2016). Antibodies were generated against three phosphopeptides containing pS356/pT357, pT363 or pS369, respectively. Affinity-purified antibodies were extensively characterized and found to be highly specific in immunoblotting for the mouse KOPR phosphorylated at S356/T357, T363 or S369 (Chen et al., 2016). U50,488H markedly enhanced staining intensity of the KOPR by immunoblotting with these three phospho-specific antibodies and the U50,488H effect was blocked by the selective KOPR antagonist norbinaltorphimine, indicating that U50,488H-induced KOPR phosphorylation at these sites is KOPR-mediated (Chen et al., 2016).

However, the regulatory mechanisms of site-specific KOPR phosphorylation are not clear yet. In this study, we focused on agonist-dependent and agonist-independent KOPR phosphorylation. Which protein kinases were involved in site-specific KOPR phosphorylation and how protein kinase-mediated KOPR phosphorylation regulated functional outputs were examined.

## Materials and methods

### Materials

Mouse neuro2A neuroblastoma (N2A) cells were purchased from the A.T.C.C. (Manassas, VA). U50,488H were provided by the National Institute on Drug Abuse (Bethesda, MD). Horseradish peroxidase (HRP)-conjugated goat anti-rabbit or anti-mouse IgG antibodies were purchased from Jackson ImmunoResearch Laboratories (West Grove, PA). Minimal essential medium (MEM), blasticidin and Lipofectamine^TM^ 2000 were purchased from Invitrogen/Thermo Fisher Scientific (Waltham, MA). Fetal bovine serum was purchased from Atlanta Biochemicals (Flowery Branch, GA). Protease inhibitor tablets and enhanced chemiluminescence (ECL) reagent Supersignal West Pico were purchased from Pierce/Thermo Fisher Scientific (Waltham, MA). Nickel-nitrilotriacetate (Ni-NTA) agarose was purchased from Qiagen (Hilden, Germany). Dodecyl-β-D-maltoside (DDM), Go6976 and PVDF membranes were purchased from EMD Millipore (Billerica, MA). Chelerythrine chloride (CHL) was from Enzo Life science (Farmingdale, NY). [^3^H]diprenorphine (specific activity 37 Ci/mmole) was purchased from PerkinElmer (Waltham, MA). Dynorphin A(1-17) was from Multiple Peptide System (SD, CA). Antibodies against total ERK1/2 (#4696) and phosphorylated ERK1/2 (#4370) were from Cell Signaling Technology (Danvers, MA). Secondary antibodies IRDye 800CW goat anti-rabbit IgG and IRDye680 goat anti-mouse IgG were from LI-COR Bioscience (Lincoln, NE). Takeda compound 101 (Compound 101; 3-[[[4- methyl-5-(4-pyridyl)-4H-1,2,4-triazole-3-yl] methyl] amino]-N-[2-(trifuoromethyl) benzyl] benzamidehydrochloride) (Lowe et al., 2015)was from Hello Bio Inc. (Princeton, NJ). Antibodies against β-arrestin1 and β-arrestin2 were generous gifts from Dr. Jeffrey Benovic of Thomas Jefferson University (Philadelphia, PA). GRK5 cDNA in pcDNA3 was kindly given by Dr. Walter Koch of Temple University Lewis Katz School of Medicine (Philadelphia, PA).

The rabbit antibodies against mKOPR (PA847) and phospho-specific KOPR were custom-generated by Covance (Conshohocken, PA) and purified in our laboratory (Chen et al., 2016). Antigens used were mKOPR (352-360) with pS356/pT357, mKOPR(357- 367) with pT363, mKOPR(365-373) with pS369, respectively.

siRNAs targeting GRK2 (sc-35513), GRK3 (sc-35515), β-arrestin1 (sc-29742), β- arrestin2 (sc-29743), PKCα (sc-36244) and PKCε (sc-36250) and antibodies against GRK2 (sc-562), GRK3 (sc-563) were purchased from Santa Cruz Biotechnology (Santa Cruz, CA). Polymerase chain reaction (PCR) primers for PKC isoforms (PKCα, PKCβ, PKCγ, PKCζ, PKCι/λ, PKCδ, PKCε, PKCθ and PKCη) were also purchased from Santa Cruz Biotechnology. GRK4/5/6 antibody (#05-466) was purchased from Millipore (Billerica, MA). siRNAs against GRK5 and GRK6 and TRIzol® Reagent were bought from Ambion by Life Technologies (Carlsbad, CA). The sense sequences of all the siRNAs used are listed in the Supplementary Materials Table S1. GF109203X, penicillin/streptomycin, pyruvic acid, NaHCO_3,_ phorbol-12-myristate-13-acetate (PMA), tetrasodium pyrophosphate, glycerophosphate disodium, NaF and orthovanadate (Na_3_VO_4_) and all other chemicals were purchased from Sigma-Aldrich (St. Louis, MO).

### Cell Culture

The mKOPR tagged with FLAG at the N-terminus and 6×His-tag at the C-terminus (FmK6H) was cloned into the vector pcDNA6. FmK6H was stably transfected into N2A cells and stable clonal N2A-FmK6H cells were selected (Chen et al., 2016). Cells were grown in 10% fetal bovine serum, 2.2 g L^-1^ NaHCO_3_, 110 mg L^-1^ pyruvic acid, 1 µg mL^-1^ blasticidin, MEM and 100 units mL^-1^ penicillin/streptomycin in a humidified atmosphere with 5% CO_2_, 95% air at 37°C.

### Drug treatment

#### Effects of U50,488H on KOPR phosphorylation: time course and dose-response relationship

Cells at 80% confluency were incubated in serum-free medium for at least 1 hr. For time-course experiments, cells were treated with vehicle or 0.1 µM U50,488H for different time intervals (0, 0.5, 1, 2, 5, 10 min) at 37°C. For concentration-response relationship, cells were treated with vehicle or different concentrations of U50,488H (0.01, 0.1, 0.3, 1, 3, 10 µM) for 2 min at 37°C.

#### Roles of GRKs in U50,488H-promoted KOPR phosphorylation

For knockdown of GRK isoforms, 5×10^5^ cells/ well in 6-well plates were left untransfected or transfected with control siRNA (siCtrl) or siRNA targeting GRK2, GRK3, GRKs2+3 or GRK6 (75 pmole siRNA) using Lipofectamine 2000. Because GRK5 was not detected in N2A cells, effects of both overexpression and overexpression/ knockdown of GRK5 were examined. Cells were co-transfected with pcDNA3/ siCtrl, GRK5 cDNA/ siCtrl, pcDNA3/ GRK5 siRNA and GRK5 cDNA/ GRK5 siRNA (1 µg cDNA/100 pmole siRNA) using Lipofectamine 2000. Two days after transfection, cells were treated with vehicle or 3 µM U50,488H for 2 min at 37°C. Some cells were pretreated with compound 101 (3 and 30 µM) [concentrations based on (Lowe et al., 2015)] for 30 min and then incubated with vehicle or 3 µM U50,488H for 2 min.

#### Effects of protein kinase C (PKC) activation on KOPR phosphorylation

Cells (∼10^7^) were incubated in serum-free medium for at least 1 hr, and then treated with different concentrations (0.001, 0.01, 0.1, 1 and 10 µM) of PMA, a PKC activator, for 30 min or treated with 0.1 µM PMA for different time points (0, 1, 5, 10 and 30 min) at 37°C. For comparison, 3 µM U50,488H was used for the same time intervals.

#### Effects of PKC inhibition on U50,488H-induced KOPR phosphorylation

Cells (∼10^7^) were incubated with serum-free medium for 1 hr followed by incubation with vehicle or a PKC inhibitor [4 µM GF109203X, 1 µM chelerythrine chloride (CHL) or 100 nM Go6976] [concentrations based on (Herbert et al., 1990; Jacobson et al., 1995; Martiny-Baron et al., 1993)] for 30 min and then were treated with vehicle or 3 µM U50,488H for 2 min at 37°C. Some cells were transfected with control siRNA (siCtrl) or siRNA targeting PKCα or PKCε (75 pmole siRNA) using Lipofectamine 2000 and the 2 days later, cells were treated 3 µM U50,488H for 2 min.

### Receptor purification and western blot

Receptor purification and subsequent immunoblotting were performed according to a simplified procedure of Chen et al. (Chen et al., 2016). Briefly, after treatment, cells were lysed with lysis buffer [1% DDM in 20 mM Tris-HCl, pH 7.4, containing phosphatase inhibitors (20 mM tetrasodium pyrophosphate, 20 mM glycerophosphate disodium, 20 mM NaF and 2 mM Na_3_VO_4_) and protease inhibitors] and centrifuged. The supernatant was incubated with Ni-NTA agarose for 1 hr and then washed with wash buffer (0.1% DDM, 20 mM imidazole and 0.5N NaCl in 20 mM Tris-HCl, pH 7.4 containing phosphatase inhibitors at 1/10 concentration). Samples were eluted with elution buffer (0.3 M imidazole, 66 mM Tris-free base, 0.3 M NaCl and 0.05% DDM), mixed with 5X sample loading buffer (10% SDS, 0.25 M Tris, pH 7.4, 50% glycerol and 0.01% Bromophenol Blue, 25 µl), resolved with 8% SDS-PAGE and transferred onto PVDF membranes. Membranes were incubated with phospho-specific KOPR antibodies (1 µg ml^-1^, pS356/pT357, pT363 and pS369) overnight followed by HRP-conjugated goat anti-rabbit IgG antibodies and then reacted with ECL reagents. Images were captured with LAS 1000 plus Image Analyzer (Fuji Photo Film Co. Ltd., Tokyo, Japan). Membranes were stripped and re-blotted for total mouse KOPR (anti-PA847, 10,000^-1^). Intensities of phospho-KOPR were normalized against that of the total KOPR in the same lane. Unless indicated otherwise, a U50,488H response (usually the maximal response) was designated as 100% and others were normalized against it.

### Reverse transcription- polymerase chain reaction (RT-PCR)

Cells (∼10^7^) were solubilized with 1 ml Trizol reagent followed by RNA extraction and precipitation for RT-PCR. The following PKC isoforms were examined for their presence in N2A cells: conventional PKCs (PKCα, PKCβ and PKCγ), atypical PKCs (PKCζ and PKCι/λ), and novel PKCs (PKCδ, PKCε, PKCθ and PKCη). RT-PCR was performed according to the semi-quantitative nested RT-PCR protocol of the manufacturer (Santa Cruz). No-primer reaction was used as control (RT-ctrl). The PCR cycle was as follows: initial denature at 95°C for 30 sec followed by 35 cycles (95°C, 45 sec; 55°C, 45 sec; 72°C, 1 min) and then the final elongation at 72°C for 10 min. PCR products were resolved with 1% agarose gel electrophoresis and the image was captured with the LAS 1000 plus Image Analyzer.

### KOPR internalization

Some cells were treated with 0.1 µM PMA for 30 min. Some other cells were incubated with a PKC inhibitor (4 µM GF109203X or 1 µM CHL) for 30 min followed by 3 µM U50,488H for 30 min. Cells were quickly suspended in PBS containing 1mM EDTA and then washed three times for 10 min with 0.1% BSA in PBS at 4°C to remove PMA or U50,488H. KOPR internalization was determined by radioligand binding to KOPR in intact cells as described previously (Li et al., 1999). Receptor binding was carried out 1 nM [^3^H]diprenorphine. using Specific binding to total receptors was defined in the presence of naloxone (10 µM), whereas specific binding to cell surface receptors was determined in the presence of dynorphin A-(1-17) (1 µM). Thus, the intracellular receptor pool can be calculated by subtracting cell surface receptor binding from total receptor binding. The internalized receptors were calculated as (intracellular receptor pool of U50,488H-treated group - intracellular receptor pool of control group) / intracellular receptor pool of control group.

### ERK1/2 activation by U50,488H

For time course study, cells were treated with 3 µM U50,488H at 37°C for different durations (0, 1, 2, 5, 10, 20, 30 min and 1, 2, 3, 4 hr). For involvement of G proteins, cells were treated with or without PTX (200 ng ml^-1^, 2 hr, 37°C) and subsequently with 3 µM U50,488H or vehicle for 5 min or 1 hr. For the roles of GRKs and β-arrestins, cells were transfected with siRNA control or siRNA against GRK2+3, GRK6, β-arrestin1 or β-arrestin2 for 2 days as described above and then treated with 3 µM U50,488H for 5 min or 1 hr at 37°C.

All treated cells were lysed with 2x Laemmli sample buffer (4% SDS, 100 mM DTT, 10% glycerol, 62.5 mM Tris, 0.01% bromophenol blue, pH6.8). Cell lysate was subject to 8% SDS-PAGE and resolved protein bands were then transferred onto nitrocellulose membranes. Membranes were incubated with primary antibodies against β- arrestin1 (1:5000), β-arrestin2 (1:6000), GRK2 (1:1000), GRK3 (1:1000), GRK4/5/6 (1:2000) or pERK1/2 (1:3000) and total ERK1/2 (1:3000) overnight at 4°C followed by goat anti-mouse IgG conjugated with infrared dye 680 or goat anti-rabbit IgG conjugated with infrared dye 800CW (1:10,000). Images were captured with the Odyssey Infrared Imaging System (LI-COR Biosciences, Lincoln, NE).

### Data Analysis and Statistics

Data were graphed and analyzed with Graph Pad Prism5.01 (La Jolla, CA). One-way ANOVA was used to analyze one-factor experiment, followed by Newman-Keuls *post-hoc* test. Two-way ANOVA was used to analyze two-factor experiments followed by Bonferroni or Newman-Keuls *post-hoc* test.

## Results

### Time course of U50,488H-induced KOPR phosphorylation

N2A-FmK6H cells were treated with 0.1 µM U50,488H for 0.5, 1, 2, 5 and 10 min. U50,488H very rapidly induced KOPR phosphorylation at S356/T357, T363 and S369, reaching a plateau in 30 sec, 1 min and 2 min, respectively (Fig. 1A). Thus, for the subsequent experiments, 2-min incubation time with U50,488H was chosen. In addition, S356/T357 showed basal phosphorylation without U50,488H treatment, similar to our previous finding (Chen et al., 2016).

**Figure 1.**
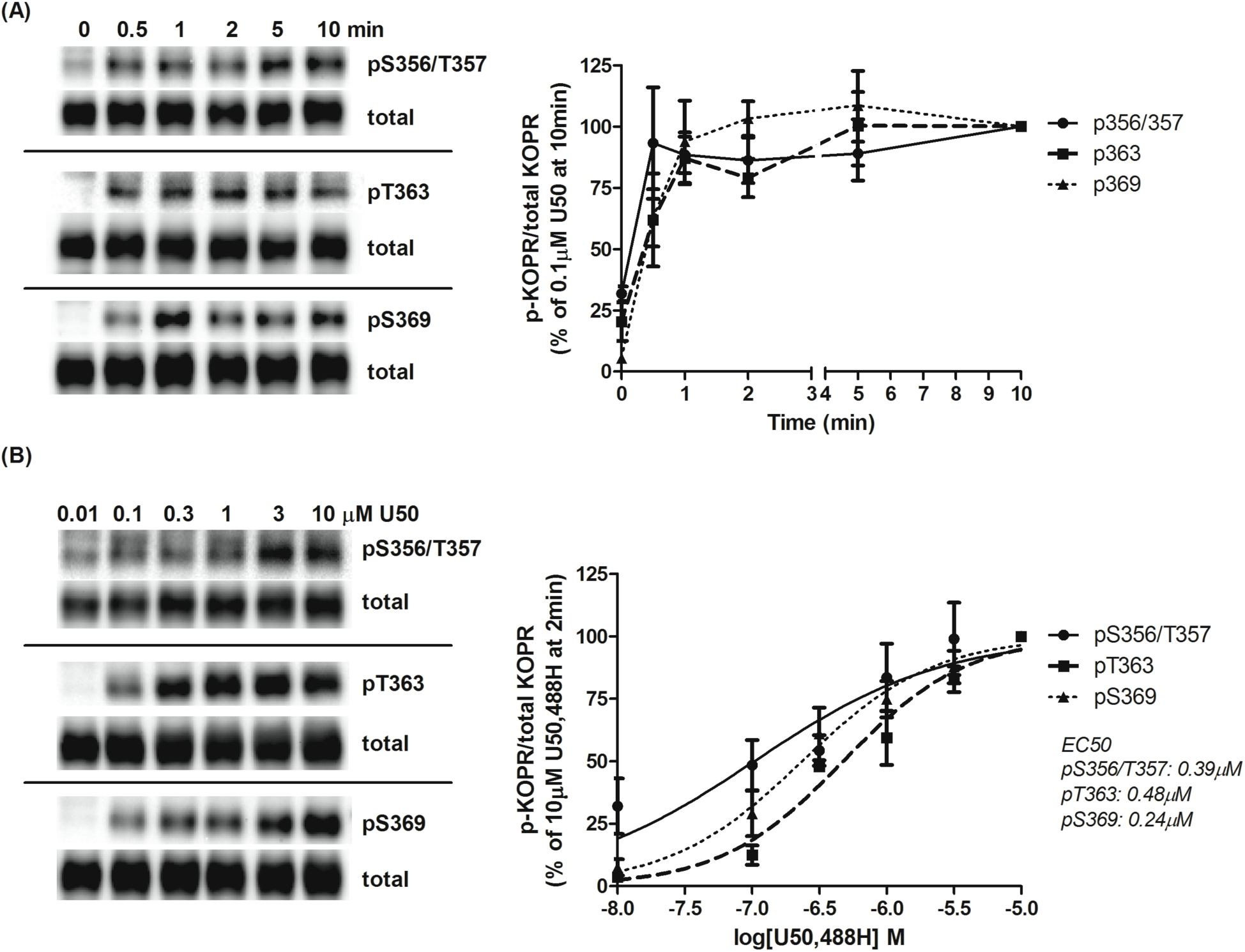
Time course and dose-response relationship of U50,488H-induced KOPR phosphorylation. N2A-FmK6H cells were treated with (A) 0.1 µM U50,488H for different durations (0, 0.5, 1, 2, 5 and 10 min) or (B) different concentrations of U50,488H (0.01, 0.1, 0.3, 1, 3 and 10 µM) for 2 min at 37°C. Cells were lysed with lysis buffer and then receptors were affinity-purified using Ni-NTA agarose beads. Eluted samples were subject to western blot using the three phospho-KOPR specific antibodies. Membranes were stripped and blotted again for total KOPR. See methods for details. One representative blot of each phospho-KOPR antibody is shown. Staining intensity was quantified and normalized against that of 10-min treatment (A) or 10 µM U50,488H (B). Each value is mean ± SEM (n=3-7).

### Dose-response relationship of U50,488H-induced KOPR phosphorylation

Cells were treated with 0.01, 0.1, 0.3, 1, 3 and 10 µM U50,488H for 2 min. U50,488H promoted KOPR phosphorylation in a dose-dependent manner at each residue (Fig. 1B). The pEC_50_ values (95% confidence limits, CI) for phosphorylation at S356/T357, T363 and S369 were determined to be 6.40 (95% CI, 6.07-6.76), 6.32 (95% CI, 5.69-6.95) and 6.63 (95% CI, 6.46-6.79), respectively. The means of EC_50_ values for S356/T357, T363 and S369 phosphorylation were 0.39 µM, 0.48 µM and 0.24 µM, respectively. There was no significant difference among the three EC_50_ values (one-way ANOVA followed by Newman-Keuls *post-hoc* test, p=0.99). Based on these results, 3 µM U50,488H was chosen for the following experiments.

### Roles of Gi/o proteins in U50,488H-indcued KOPR phosphorylation

Cells were pretreated with pertussis toxin (PTX) followed by U50,488H incubation. PTX significantly reduced U50,488H-induced KOPR phosphorylation at S356/T357, T363 and S369 by 81 ± 8%, 52 ± 6% and 62 ± 5%, respectively (Fig. 2). The reduction in S356/T357 phosphorylation was significantly higher than T363 phosphorylation (one-way ANOVA followed by Newman-Keuls *post-hoc* test, p<0.05). The results indicate activation of G_i/o_ proteins is involved in KOPR phosphorylation at each phosphosite and pS356/T357 was more highly regulated by Gi/o proteins than other residues.

**Figure 2.**
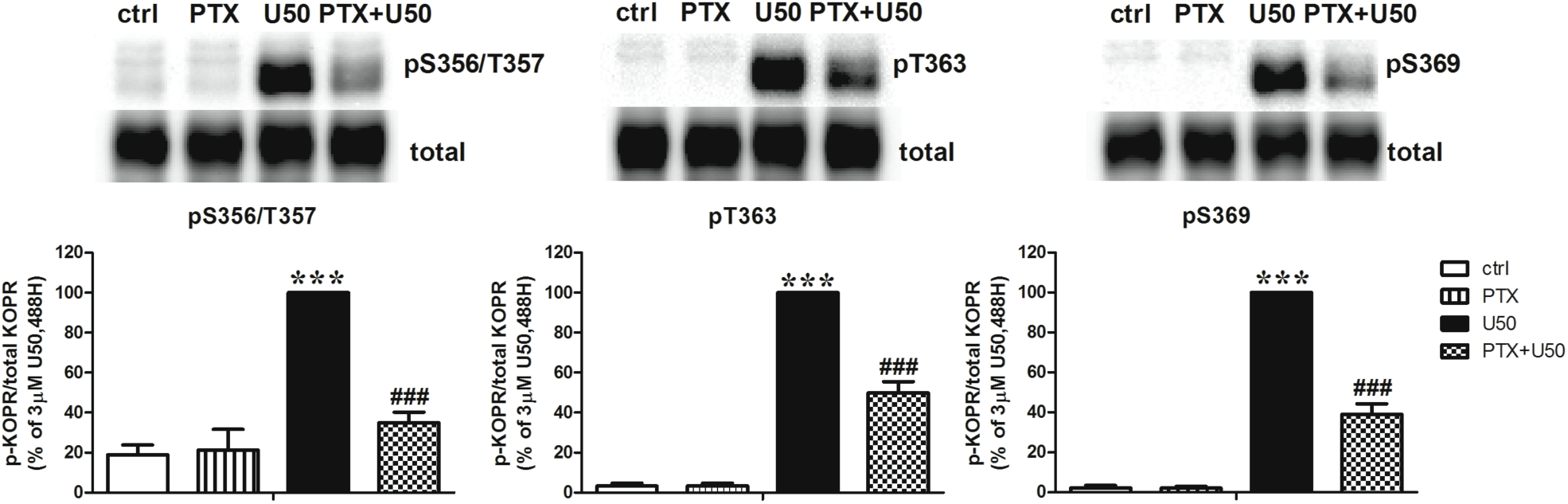
Effects of PTX on U50,488H-induced KOPR phosphorylation. Cells were pretreated with 200 ng/ml PTX for 2 hr followed by a 2-min incubation with 3 µM U50,488H. Experiments were performed as described in Fig. 1 legend. The data shown are the mean ± S.E.M (n=3) and were analyzed by two-way ANOVA followed by Newman-Keuls *post-hoc* test (***: *p* <0.001, compared to control group; ^###^: *p* <0.001, compared to U50,488H-treated group).

### Roles of GRKs in U50,488H-induced KOPR phosphorylation

GRKs are major protein kinases involved in agonist-induced GPCR phosphorylation. We found that N2A cells endogenously express GRK2, GRK3 and GRK6, but not GRK5, by western blot with GRK2, GRK3 and pan GRKs4/5/6 antibodies (see Fig. 3).

**Figure 3.**
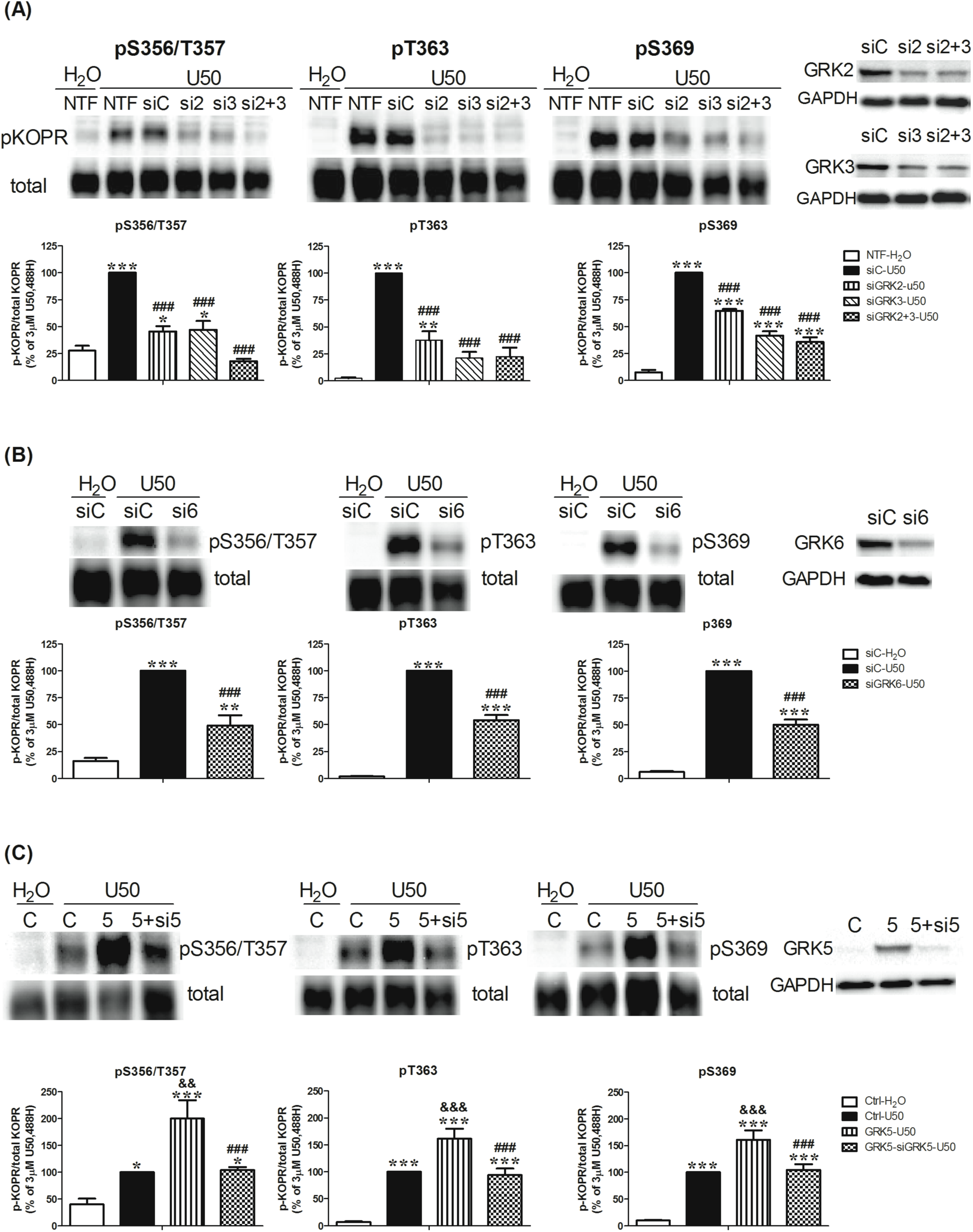
Involvement of GRKs in U50,488-induced KOPR phosphorylation. **(A, B) Effects of knockdown of GRK2, GRK3 and GRK6 on U50,488H-induced KOPR phosphorylation** Cells were left untransfected (NTF) or transfected with control siRNA (siC) or siRNAs targeting GRK2 (si2), GRK3 (si3), GRK2+3 (si2+3) or GRK6 (si6). Two days later, cells were treated with vehicle or 3 µM U50,488H for 2 min. Data were normalized against that of U50,488H-treated siRNA control group (siC-U50) and shown as the mean ± S.E.M (n=3-6). Knockdown of GRK2, GRK3 and GRK6 was verified by western blot with specific antibodies. **(C) Effects of GRK5 expression and expression + knockdown on U50,488H-induced KOPR phosphorylation** Cells were co-transfected with the vector pcDNA3 and control siRNA (C), GRK5 cDNA and control siRNA (5), and GRK5 cDNA and GRK5 siRNA (5+si5) with Lipofectamine 2000. Two days later, cells were treated with vehicle or 3 µM U50,488H for 2 min. The results were normalized against that of U50,488H-treated vector control group (Ctrl-U50) and shown the mean ± S.E.M (n=6-8). The expression of GRK5 and knockdown of GRK5 were validated by western blot. Data were analyzed with one-way ANOVA followed by Newman-Keuls *post-hoc* test [*: *p* <0.05, **: *p* <0.01, ***: *p* <0.001, compared to vehicle control group (siC-H_2_O) or H_2_O-treated control group (Ctrl-H_2_O); ^###^: *p* <0.001, compared to U50,488H-treated siRNA control group (siC-U50) or U50,488H-treated GRK5 overexpression group (GRK5-U50); ^&&^: *p* <0.01, ^&&&^: *p* <0.001, compared to U50,488H-treated vector control group (Ctrl-U50)].

Knockdown of GRK2 or GRK3 decreased U50,488H-promoted phosphorylation at S356/T357 by 77 ± 12% and 76 ± 16%, respectively, whereas knockdown of both totally blocked phosphorylation (115 ± 11%). All the values are mean ± SEM (n=3-6). For T363, knockdown of GRK2 reduced phosphorylation by 63 ± 9%, while knockdown of GRK3 and GRKs2/3 reduced by 80 ± 5% and 79 ± 8 %, respectively. For S369, knockdown of GRK2 inhibited phosphorylation by 38 ± 3%, whereas knockdown of GRK3 and GRKs2/3 reduced by 62 ± 4% and 68 ± 4%, respectively (Fig. 3A). The control siRNA did not affect U50,488H-promoted phosphorylation at S356/T357, T363 and S369, compared to no transfection (NTF). Therefore, NTF was omitted in the subsequent experiments involving siRNA. Knockdown of GRK6 reduced U50,488H- caused KOPR phosphorylation at S356/T357, T363 and S369 by 60 ± 11%, 47 ± 5% and 53 ± 5%, respectively (Fig. 3B). These results indicate that GRK2, GRK3 and GRK6 are involved in KOPR phosphorylation at all four residues upon U50,488H stimulation. GRK2 alone or GRK2+3 shows higher involvement at S356/T357 than T363 and S369 (Fig. 7).

GRK5 was reported to participate in agonist-induced KOPR desensitization (Appleyard et al., 1999), suggesting that GRK5 is involved in KOPR phosphorylation. Since N2A cells do not express GRK5 endogenously, we examined the role of GRK5 by its expression and expression + knockdown. Expression of GRK5 promoted higher levels of KOPR phosphorylation at S356/T357, T363 and S369 by 2-, 1.6- and 1.6-fold, respectively, compared with the vector control. Co-transfection of GRK5 and GRK5 siRNA blocked GRK5 expression-caused KOPR phosphorylation at S356/T357, T363 and S369 by 64 ± 9%, 42 ± 8% and 38 ± 3 % (Fig. 3C). GRK5 is more involved in phosphorylating S356/T357 than the other two residues (Fig. 7).

Knockdown efficiency for each GRK was determined (Fig. 3 left panel). GRK2 siRNA or GRK2 + 3 siRNAs reduced GRK2 by 47 ± 6% and 48 ± 4%, respectively, compared to control siRNA. Similarly, GRK3 siRNA or GRK2 + 3 siRNAs reduced GRK3 by 48 ± 8% and 47 ± 7%, respectively. GRK6 siRNA reduced GRK6 levels by 40 ± 5%. GRK5 siRNA reduced the overexpression level of GRK5 by 79 ± 3%. The specificity of each individual siRNA against GRKs 2, 3, 5 and 6 for its target vs. other GRKs was confirmed (data not shown).

We also used the GRK2/3 inhibitor compound 101 (Lowe et al., 2015) to examine the roles of GRK2 and GRK3 in U50,488H-induced KOPR phosphorylation. Compound 101 (3 and 30 µM) dose-dependently inhibited U50,488H-promoted KOPR phosphorylation at S356/T357, T363 and S369 and at 30 µM completely blocked U50,488H-induced KOPR phosphorylation at all the four residues (Supplemental Fig. S1). The results are similar to those obtained with siRNA. Compound 101 at 3 µM showed similar levels of reduction as knockdown of GRK2+3. GRK2+3 and compound 101 regulated S356/T357 more than T363 and S369 (Fig. 7). Compound 101 at 30 µM reduced basal S356/T357 phosphorylation.

Taken together, U50,488H-induced KOPR phosphorylation at S356/T357, T363 and S369 was regulated by GRK2, GRK3, GRK5 and GRK6. All four phosphoresidues are regulated concertedly by the four GRKs. GRK2, GRK2+3 and GRK5 appear to more involved in phosphorylating S356/T357 than T363 and S369 (S356/T357> T363= S369) (Fig. 7). GRK3 and GRK6 are similarly involved in phosphorylating the four residues (S356/T357= T363= S369) (Fig. 7).

### Roles of PKCs in KOPR phosphorylation

PKC was reported to be involved in homologous and heterologous phosphorylation of GPCRs, including MOPR and free fatty acid receptor4 (Burns et al., 2014; Mann et al., 2015). Activation of the KOPR by agonists has been shown to stimulate PKC (Bian et al., 2000; Bohn et al., 2000). S356, T357, T363 and S369 are also PKC consensus phosphorylation sites. Thus, whether PKC regulates agonist-independent and agonist-dependent KOPR phosphorylation was characterized.

Using RT-PCR, we have found that N2A cells endogenously express eight PKC isoforms: PKCα, PKCβ and PKCγ (conventional PKCs), PKCζ and PKCι/λ (atypical PKCs), as well as PKCδ, PKCε and a low level of PKCθ, but not PKCη (novel PKCs) (Fig 4).

**Figure 4.**
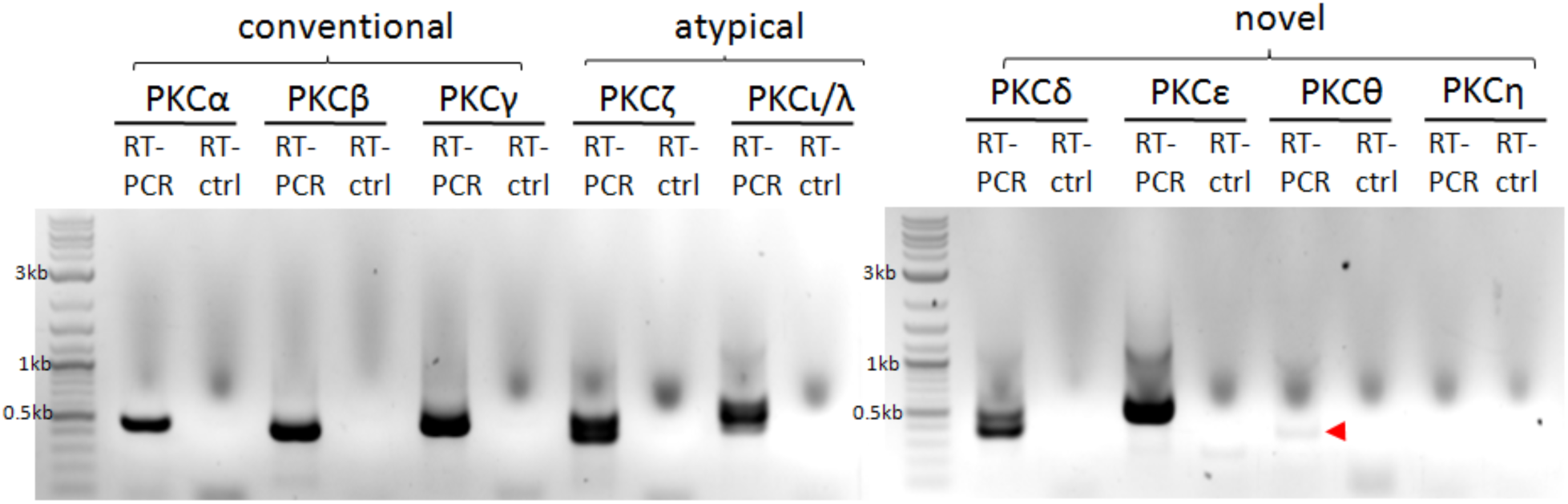
Endogenous PKC isoforms in N2A cells: detected with RT-PCR. N2A cells were solubilized with Trizol reagent and then RNA was extracted. RNA was used for RT-PCR for detection of different PKC isoforms (conventional PKCs: PKCα, PKCβ and PKCγ; atypical PKCs: PKCζ and PKCι/λ; novel PKCs: PKCδ, PKCε, PKCθ and PKCη). For the control, no PCR primers were added (RT-ctrl). The following PKC isoforms (predicted PCR product sizes) were detected: PKCα (496bp), PKCβ (428bp), PKCγ (552bp), PKCζ (428bp), PKC?/λ (456bp), PKCδ (428bp), PKCε (547bp), and PKCθ (494bp). PKCθ PCR product was barely detectable (red arrowhead). No PKCη PCR product was detected. The experiment was carried out twice with similar results.

#### Roles of PKCs in agonist-dependent KOPR phosphorylation

Knockdown of PKCα or PKCε by isoform-specific siRNA, which reduced each isoform level by 45% and 43%, respectively, did not significantly affect U50,488H- induced KOPR phosphorylation at any of the four sites (Supplemental Fig. S2). It is likely that siRNA knockdown of one isoform may lead to substitution by other PKC isoforms. Thus, PKC inhibitors were used subsequently to examine agonist-dependent KOPR phosphorylation.

The non-selective PKC inhibitor GF109203X (4 µM) reduced U50,488H promoted KOPR phosphorylation at S356/T357, T363 and S369 by about 100 ± 1%, 59 ± 10% and 27 ± 12%, respectively (Fig. 5A). The extents of reduction by GF109203X were significantly different among the residues (Fig. 7). In addition, GF109203X lowered the basal phosphorylation level of S356/T357, but not T363 and S369 (Fig. 5A). Another non-selective PKC inhibitor chelerythrine (CHL) (1 µM) significantly decreased U50,488H-promoted KOPR phosphorylation at T363 and S369 by 58 ± 10% and 61 ± 5%, respectively, but had no effect at S356/T357 (16 ± 12%), without affecting basal phosphorylation (Fig. 5B). The selective PKCα and PKCβ1 inhibitor Go6976 (100 nM) reduced U50,488H-induced KOPR phosphorylation at S356/T357 and S369 by 36 ± 18% and 22 ± 12%, respectively, but not at T363 (-7 ± 11%), with no effect on basal phosphorylation (Fig. 5C). These results indicate different PKC isoforms contribute to U50,488H-induced KOPR phosphorylation in site-specific manner. In addition, PKC is involved in basal KOPR phosphorylation at S356/T357.

**Figure 5.**
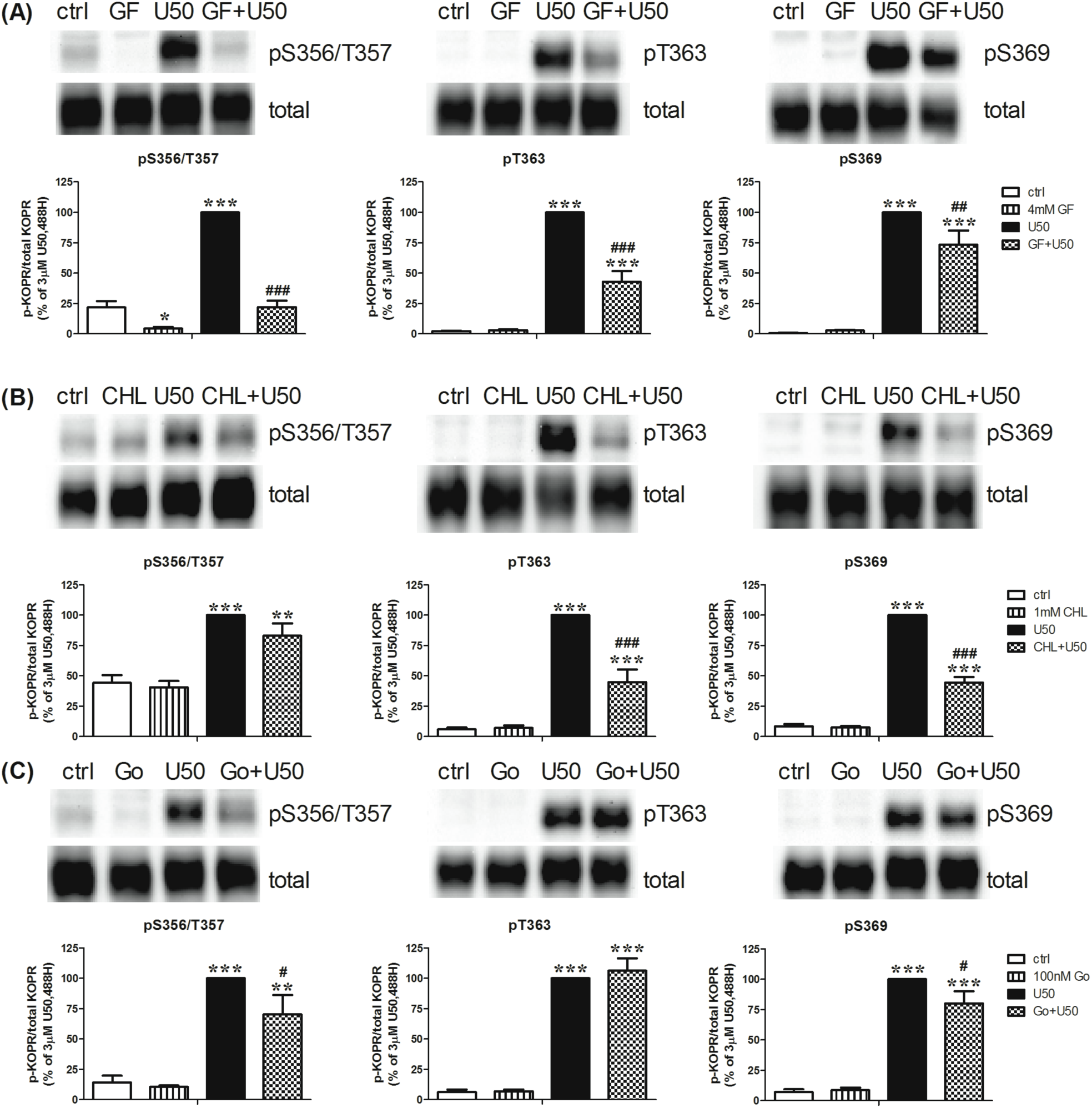
Effect of PKC inhibitors on U50,488H-induced KOPR phosphorylation at S356/T357, T363 and S369. Cells were pretreated with **(A)** the nonselective PKC inhibitor GF109203X (GF) (4 µM), **(B)** chelerythrine chloride (CHL) (1 µM) or **(C)** the selective PKC α and β1 inhibitor Go6976 (100 nM) for 30 min followed by 3 µM U50,488H or vehicle for 2 min. Each point is shown as the mean ± S.E.M (n=3-6). Data were analyzed by two-way ANOVA followed by Newman-Keuls *post-hoc* test (*: *p* <0.05, **: *p* <0.01, ***: *p* <0.001, compared to control group; ^#^: *p* <0.05, ^##^: *p* <0.01, ^###^: *p* <0.001, compared to U50,488H- treated group).

#### Activation of PKCs increased agonist-independent KOPR phosphorylation

Whether PKC activation promoted agonist-independent KOPR phosphorylation was studied using the PKC activator PMA. Incubation of cells with different concentrations of PMA (0.001, 0.01, 0.1, 1 and 10 µM) for 30 min enhanced KOPR phosphorylation at S356/T357, T363 and S369 in a dose-dependent manner. The pEC_50_ (95% confidence limits, CI) of PMA in promoting phosphorylation at S356/T357, T363 and S369 were determined to be: 8.18 (95% CI 7.85-8.5), 7.50 (95% CI 7.00-8.03) and 7.72 (95% CI 7.55-7.89), respectively. The mean of EC_50_ values of S356/T357, T363 and S369 phosphorylation were 6.6 nM, 31.8 nM and 19 nM, respectively (Fig. 6A). The EC_50_ values were significantly different between S356/T357 and T363 (p=0.045, one-way ANOVA followed by Newman-Keuls *post-hoc* test). Thus, PKC activation promoted agonist-independent KOPR-phosphorylation and PMA had lower potency in phosphorylating T363. Based on these results, 0.1 µM PMA was used in following experiments.

**Figure 6.**
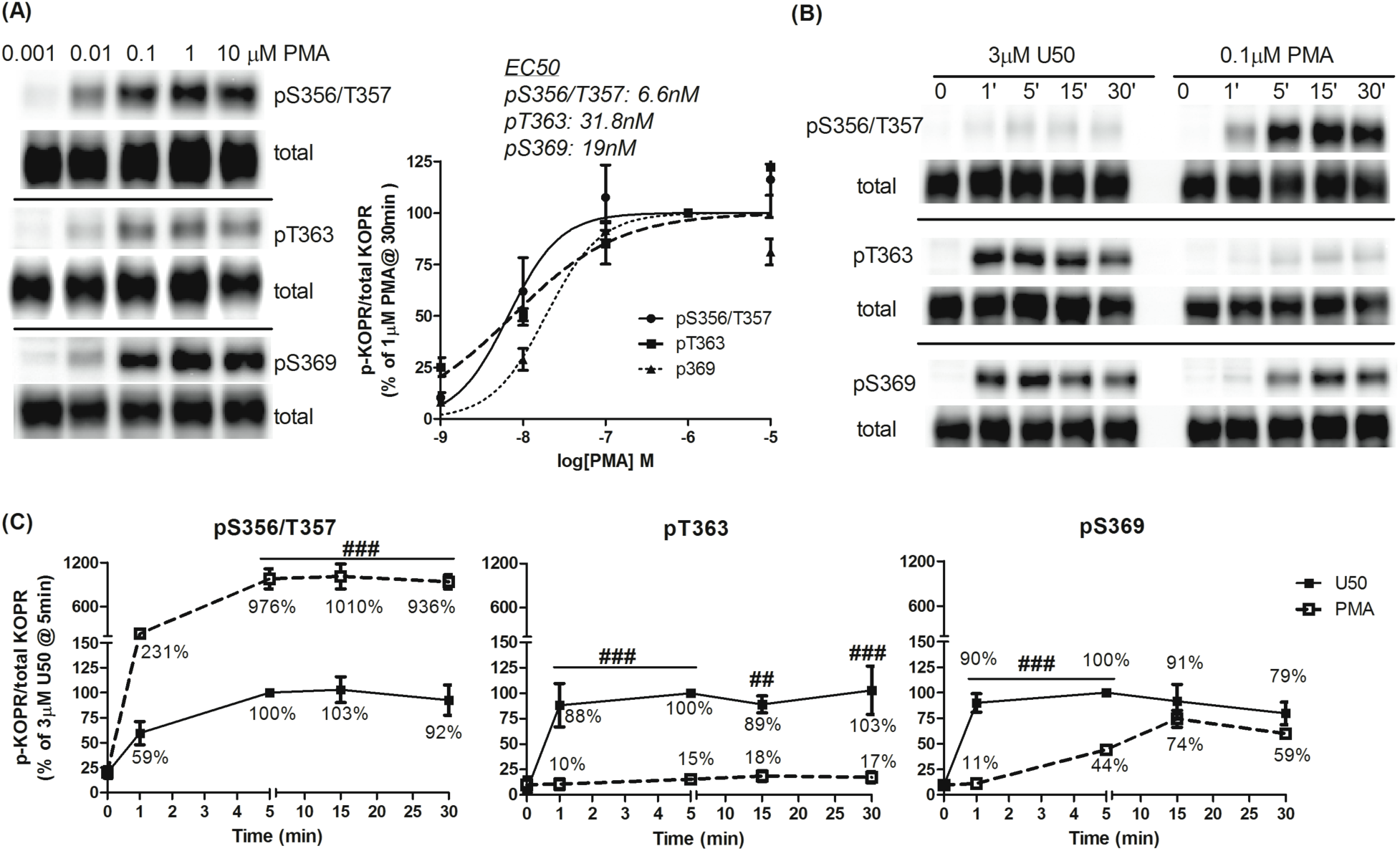
Effect of PKC activation on KOPR phosphorylation. **(A) Dose-response relationship of PMA-induced KOPR phosphorylation:** Cells were treated with different concentrations of PMA (0.001, 0.01, 0.1, 1 and 10 µM) for 30 min. **(B, C) Time course of PMA-induced KOPR phosphorylation and comparison with that of U50,488H**: Cells were incubated with 3 µM U50,488H or 0.1 µM PMA for 0, 1, 5, 15 and 30 min. In (A), staining intensity was normalized against that of 1 ?M PMA and each value is the mean ± S.E.M (n=3-4). The mean EC_50_ values of PMA in promoting phosphorylation at S356/T357, T363 and S369 are shown. In **(C)**, staining intensity in (B) was normalized against that of U50,488H at 5 min. Data were shown as the mean ± S.E.M (n=3-4) and were analyzed by two-way ANOVA followed by Bonferroni *post-hoc* test (^##^: *p* <0.01, ^###^: *p* <0.001, compared to U50,488H-treated group).

The time course and levels of PMA-promoted KOPR phosphorylation were examined and compared with those of U50,488H. Cells were treated with PMA (0.1 µM) or U50,488H (3 µM) for different durations (0, 1, 5, 15 and 30 min). As shown in Figs. 6B and 6C, the time courses were different between PMA and U50,488H. PMA displayed a slower time course at S369 than U50,488H and similar phosphorylation rates at S356/T357 as U50,488H. For S356/T357, PMA promoted a much higher plateau phosphorylation level than U50,488H (∼10 fold). For T363, U50,488H induced phosphorylation to greater extents than PMA (∼7 fold). For S369, U50,488H and PMA caused similar maximal phosphorylation levels. The results indicate that U50,488H and PMA showed different KOPR phosphorylation patterns and time courses.

U50,488H-induced KOPR phosphorylation at S356/T357 in Fig. 6B showed lower signals than in Fig. 1A. Since PMA-induced KOPR phosphorylation at S356/T357 yielded very high signals (Fig. 6B), the U50,488H-induced pS356/T357 signals in Fig. 6B were captured with shorter exposure time to prevent the PMA signals from being over-exposed. In addition, PMA-induced KOPR phosphorylation at pT363 in Fig. 6B showed weaker signal than that in Fig. 6A (see the signals at 0.1 μM, 30 min) because they were captured with shorter exposure time to prevent the U50,488H signals in Fig. 6B from overexposure.

### Phosphorylation patterns: regulation of U50,488H-induced site-specific phosphorylation by distinct protein kinases

Figure 7 summarizes the involvement various kinases in U50,488H-promoted phosphorylation of the four residues in the KOPR. Different residues were regulated to different degrees by multiple protein kinases. For GRKs, although the four GRK isoforms regulated all the phosphoresidues on KOPR, each had different degrees of involvement in phosphorylating the four sites. GRK2, GRK2+3 and GRK5 phosphorylated S356/T357 more than T363 and S369 (S356/T357> T363= S369). GRK3 and GRK6 phosphorylated all the residues to similar extents (S356/T357= T363= S369). For PKC, GF109203X suppressed U50,488H-induced phosphorylation in the degrees of S356/T357>> pT363> pS369. CHL inhibited phosphorylation of T363 and S369 similarly, but had no effect on phosphorylation of S356/T357. Go6976 suppressed KOPR phosphorylation at S356/T357 and S369 (S356/T357= S369), but not at T363. Whether differences in protein kinase-mediated KOPR phosphorylation lead to diverse functional outputs was examined below.

**Figure 7.**
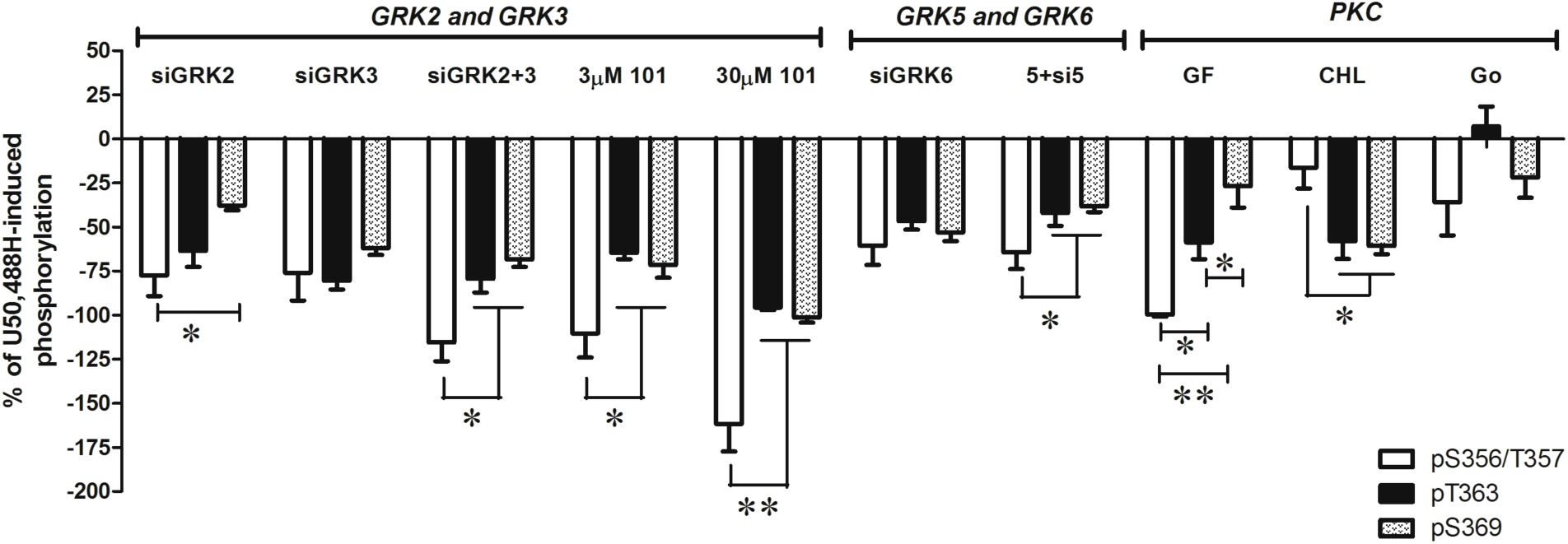
Regulation of U50,488H-KOPR phosphorylation at S356/T357, T363 and S369 by protein kinases. This figure summarizes and compares % reduction by inhibitors and GRK siRNAs of U50,488H-induced KOPR phosphorylation at all the four phosphosites. The value was calculated according to the equation: % reduction = {[(inhibitor or siRNA+ U50,488H) - U50,488H] / U50,488H}×100%. Data were analyzed by one-way ANOVA followed by Newman-Keuls *post-hoc* test (*: *p* <0.05, **: *p* <0.01).

### Effects of protein kinase-mediated KOPR phosphorylation on cellular functions

GPCR phosphorylation is the initial step in desensitizing G protein-mediated signaling as well as in initiating β-arrestin-dependent signaling and receptor internalization. Thus, we examined the effect of protein kinase-mediated KOPR phosphorylation on receptor internalization (Fig. 8) and the downstream signaling (Fig. 9).

**Figure 8.**
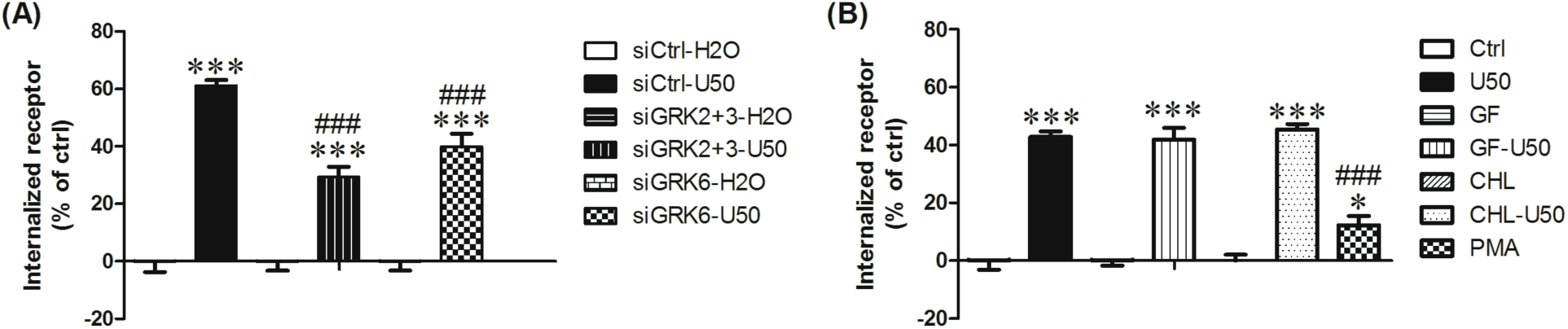
Roles of GRKs and PKC in KOPR internalization. **(A) GRKs mediated U50,488H-induced KOPR internalization** Cells were transfected with control siRNA (siCtrl) or siRNAs targeting GRK2+3 (siGRK2+3) or GRK6 (siGRK6). Two days later, cells were treated with vehicle or 3 µM U50,488H for 30 min. Cells were collected with 1 mM EDTA in PBS and then washed 3 times with 0.1% BSA in PBS for 10 min at 4°C. Receptor internalization was determined using [^3^H]diprenorphine binding to total and cell surface KOPR in cells as described in the Materials and Methods. Internalized receptors are presented as the percentage of U50,488H-treated group over water-treated group. The data were shown the mean ± S.E.M (n=3) and analyzed with one-way ANOVA followed by Newman-Keuls *post-hoc* test [***: *p* <0.001, compared to vehicle control group (siCtrl-H_2_O); ^###^: *p* <0.001, compared to U50,488H-treated siRNA control group (siCtrl-U50)]. **(B) PKC promoted agonist-independent KOPR internalization, but was not involved in U50,488H-dependent KOPR internalization** Cells were treated with 4 µM GF109203X (GF) or 1 µM chelerythrine chloride (CHL) for 30 min and then treated with water or 3 µM U50,488H for 30 min. Some cells were treated with 0.1 µM PMA for 30 min. Cells were treated as described above. Results shown are the mean ± S.E.M (n=3). Data for PKC inhibitor-treated groups were analyzed by two-way ANOVA followed by Bonferroni *post-hoc* test (***: *p* <0.001, compared to control-treated group). The data for PMA were analyzed with one-way ANOVA followed by Newman-Keuls *post-hoc* test [*: *p* <0.05, ***: *p* <0.001, compared to control group (Ctrl); ^###^: *p* <0.001, compared to U50,488H-treated group (U50)].

**Figure 9.**
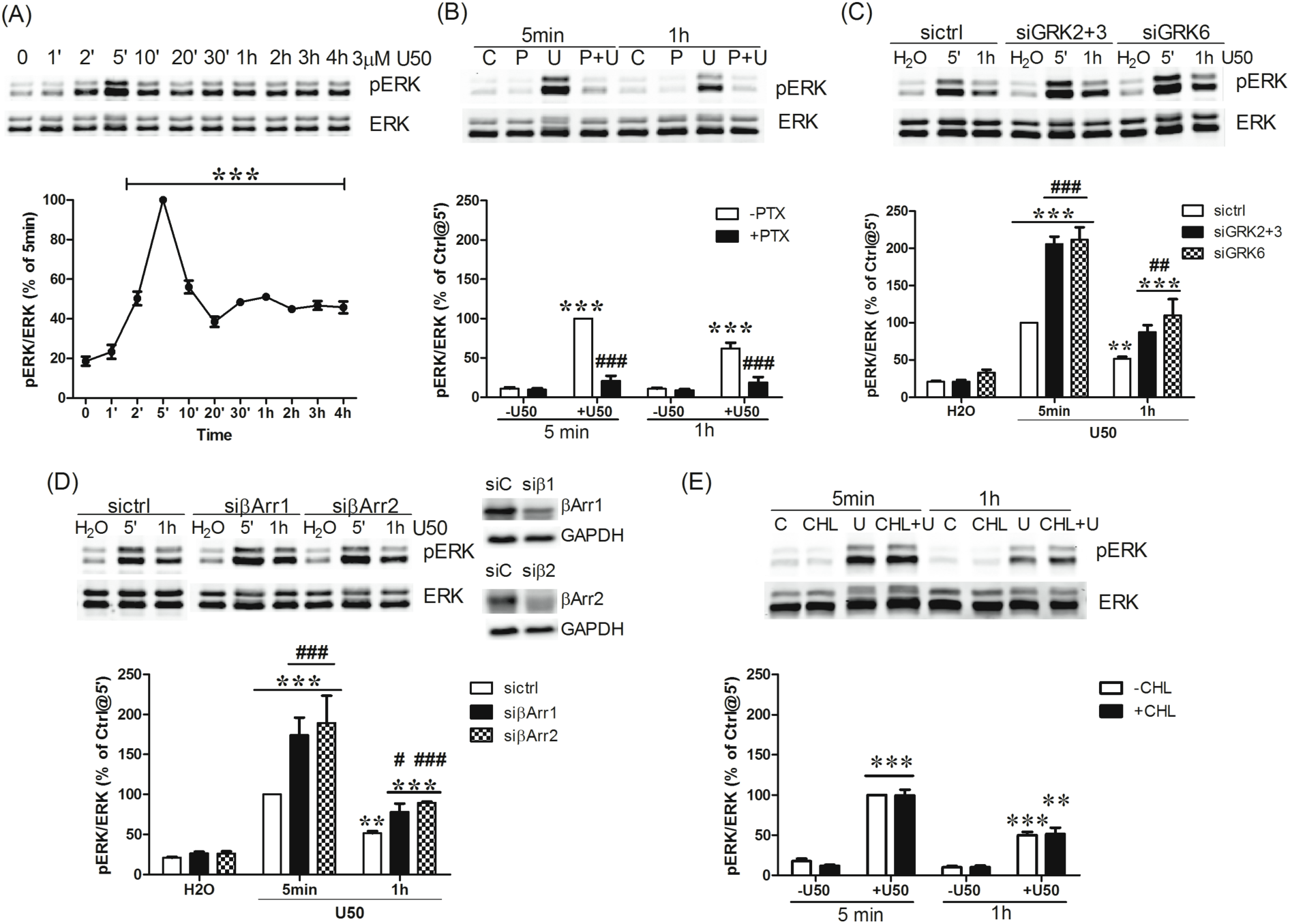
U50,488H-induced ERK1/2 phosphorylation: roles of G proteins, GRKs and β-arrestins. **(A)** Cells were treated with 3µM U50,488H for different periods of time (0, 1, 2, 5, 10, 20, 30 min and 1, 2, 3, 4 hr). **(B)** Cells were pretreated with 200 ng/ml PTX (P) for 2 hr followed by 3 µM U50,488H (U) 5 min or 1 hr. **(C, D)** Cells were transfected with control siRNA (siC) or siRNAs targeting β-arrestin1 (siβ1 or siβArr1), β-arrestin2 (siβ2 or siβArr2), GRK2+3 (si2+3) and GRK6 (si6). Two days later, cells were treated with H_2_O or 3 µM U50,488H for 5 min or 1 hr. **(E)** Cells were pretreated with 1 µM CHL for 30 min and then incubated with H_2_O or 3 µM U50,488H for 5 min or 1 hr. ERK1/2 phosphorylation was determined as described in Materials and Methods. Data shown are the mean ± S.E.M (n=3-4). Fig 9A. Data were analyzed by one-way ANOVA followed by Newman-Keuls *post-hoc* test (***: *p* <0.001, compared to time zero group). Fig 9B. Data were analyzed by two-way ANOVA followed by Bonferroni *post-hoc* test (***: *p* <0.001, compared to vehicle treated group; ^###^: *p* <0.001, compared to U50,488H- treated group). Fig 9C. Data were analyzed by two-way ANOVA followed by Bonferroni *post-hoc* test (**: *p* <0.01***: *p* <0.001, compared to water-treated group; ^##^: *p* <0.01 ^###^; *p* <0.001, compared to siCtrl-U50,488H-treated group). Fig 9D. Data were analyzed by two-way ANOVA followed by Bonferroni *post-hoc* test (**: *p* <0.01***: *p* <0.001, compared to water-treated group; ^###^: *p* <0.001, compared to sictrl-U50,488H-treated group at 5 min) and t-test for 1-hr U50,488H-treatment between sictrl and siβ-arrestin (^#^: *p* <0.05; ^###^: *p* <0.001). Fig 9E. Data were analyzed by two-way ANOVA followed by Bonferroni *post-hoc* test (**: *p* <0.01, ***: *p* <0.001, compared to vehicle treated group).

#### Effects of agonist-dependent and –independent KOPR phosphorylation on KOPR internalization

Knockdown of GRKs2/3 or GRK6 decreased U50,488H-induced KOPR internalization from ∼60% to ∼ 30% and ∼40%, respectively (Fig. 8A), indicating that GRKs2/3 and GRK6 are involved in U50,488H-induced KOPR internalization. Inhibition of PKC by GF109203X or CHL did not affect U50,488H-induced internalization (Fig. 8B). Therefore, GRK-mediated, but not PKC-mediated, KOPR phosphorylation, causes U50,488H-induced internalization. In addition, PMA induced ∼12% KOPR internalization, much lower than U50,488H (Fig. 8B).

Comparison between Fig. 8A and Fig. 8B showed that U50,488 induced internalization of 60% and 43% of cell surface KOPR, respectively. The different extents of internalization are likely due to influence of transfection on cells. Results presented in Fig. 8A involved cells with siRNAs transfection, whereas those in Fig. 8B were from cells treated with different compounds without transfection. Transfection with Lipofectamine 2000, which affects plasma membranes, may affect receptor internalization. This is a limitation of using the transfection procedures.

#### Roles of protein kinase-mediated KOPR phosphorylation in U50,488H-induced ERK1/2 signaling

KOPR activation has been shown to activate ERK1/2 (Belcheva et al., 1998; Bruchas et al., 2006). Incubation of cells with U50,488H for different durations (1, 2, 5, 10, 20, 30 min and 1, 2, 3, 4 hr) increased pERK1/2 levels with a peak at 5 min (defined as 100%), followed by a decline to about 40-50% and then sustained stimulation at 40-50% for up to 4 hr (Fig. 9A).

The roles of Gi/o proteins, GRKs and β-arrestins in U50,488H-induced ERK1/2 activation at 5 min and 1 hr were examined. PTX pretreatment blocked U50,488H- induced ERK1/2 phosphorylation at 5 min and 1 hr (Fig. 9B), indicating that both are G protein-dependent. Thus, in this N2A cell system, there is no differentiation between early phase and late phase of ERK1/2 activation as shown by McLennan et al.(McLennan et al., 2008). Knockdown of GRK2/GRK3 or GRK6 enhanced ERK1/2 activation at 5 min and 1 hr (Fig. 9C). Agonist-caused GPCR phosphorylation leads to β-arrestin recruitment, so the role of β-arrestins in U50,488H-induced ERK1/2 activation was examined. Knockdown of β-arrestin 1 or β-arrestin 2, which reduced β-arrestin 1 or β- arrestin 2 by 38 ± 5% or 45 ± 6%, respectively, increased ERK1/2 activation at 5 min and 1 hr (Fig. 9D), similar to the GRK knockdown results. These results indicate that U50,488H-induced ERK1/2 phosphorylation is G protein-, but not GRK- or β-arrestin-, dependent. Instead, GRK-mediated KOPR phosphorylation followed by β-arrestin recruitment is involved in desensitization of G protein-mediated ERK1/2 signaling.

We also examined if U50,488H-induced PKC-mediated KOPR phosphorylation regulated desensitization of G protein-mediated ERK1/2 activation. Treatment of cells with CHL (1 µM) did not affect U50,488H-induced ERK1/2 activation at 5 min and 1 hr (Fig. 9E), demonstrating that PKC is not involved. Thus, GRK-mediated, but not PKC-mediated, KOPR phosphorylation plays a role in desensitization of G protein-mediated ERK1/2 signaling.

## Discussion

Our studies have shown that Gi/o_α_ proteins and multiple protein kinases (GRKs2, 3, 5, 6 and PKC) are involved in agonist (U50,488H)-induced KOPR phosphorylation and PKC also promotes agonist-independent KOPR phosphorylation. GRK-, but not PKC-, mediated agonist-dependent KOPR phosphorylation is involved in KOPR internalization and desensitization of G protein-dependent ERK1/2 phosphorylation. PMA-induced agonist-independent KOPR phosphorylation results in lower extent of receptor internalization than U50,488H. KOPR is phosphorylated by U50,488H (GRK- and PKC-mediated) and PMA (PKC activation) in distinct phosphorylation patterns, leading to differential KOPR functional regulations (internalization and desensitization). To the best of our knowledge, this is the first demonstration that PKC and GRK6 are involved in KOPR phosphorylation and that different KOPR phosphorylation patterns cause distinct functional consequences.

### Multiple protein kinases-induced KOPR phosphorylation and the barcode hypothesis

The “barcode” hypothesis postulates that different kinases phosphorylate distinct residues of GPCRs, leading to distinct functional outcomes (Butcher et al., 2012; Nobles et al., 2011). For example, upon agonist treatment, the M3 muscarinic receptor phosphorylation is involved in desensitization of the phospholipase C pathway and receptor internalization, whereas casein kinase 1α - and casein kinase 2-mediated phosphorylation impacts on ERK1/2 and JUN kinase pathways, respectively [reviewed in (Butcher et al., 2012)]. In this study, following U50,488H treatment, GRK-mediated, but not PKC-mediated, phosphorylation is involved in U50,488H-induced internalization and desensitization of ERK1/2 signaling. PKC-activated agonist-independent KOPR phosphorylation caused lower degrees of internalization, compared to U50,488H. Thus, it is likely that fine tuning on phosphorylation of the four KOPR residues by multiple protein kinases (GRKs and PKC isoforms) via different phosphorylation patterns (residues and levels) provides a regulatory mechanism contributing to diverse cellular outcomes.

Interestingly, all four GRK isoforms (GRKs 2, 3, 5 and 6) are involved in KOPR phosphorylation at all the four residues. GRK2, GRK2+3 and GRK5 show more involvement in phosphorylating S356/T357 than T363 and S369, whereas GRK3 and GRK6 phosphorylated all the residues to similar degrees. It is noteworthy that siRNA knockdown of any of GRKs 2, 3 or 6 alone greatly decreased U50,488H-promoted KOPR phosphorylation, suggesting that concerted activities of all three endogenous GRKs are required to fully phosphorylate KOPR. Similarly, knockdown of GRKs 2/3 or GRK6 profoundly increased U50,488H-induced ERK1/2 phosphorylation and largely attenuated U50,488H-caused internalization, demonstrating that combined effects of GRKs2/3 and GRK6 are needed for full desensitization and internalization of the receptor.

GRKs have been reported to be regulated by PKC. PKC phosphorylates GRK2 and GRK5, leading to GRK2 activation, but GRK5 inhibition (Chuang et al., 1995; Pronin and Benovic, 1997; Winstel et al., 1996). N2A cells endogenously express GRK2, but not GRK5. The possibility cannot be excluded that effects of PKC inhibition are in part due to GRK2 inhibition, resulting in reduced KOPR phosphorylation. On the other hand, since compared with U50,488H, PMA induced much higher S356/T357 phosphorylation and much lower T363 phosphorylation and PMA promoted much slower S369 phosphorylation, it is unlikely that the actions of PMA were through GRK2 activation.

Different cells or tissues are likely to express different populations and levels of GRKs, PKC isoforms and other protein kinases, leading to distinct phosphorylation patterns and functional consequences. For example, U50,488H did not cause internalization of rKOPR in CHO cells (Li et al., 1999), but did so in HEK293 cells (McLaughlin et al., 2003). Therefore, GPCR phosphorylation show cell-specific and tissue-specific, leadind to recruitment of different levels of β-arrestins, activation of different downstream signal pathways and ultimately unique physiological and pharmacological responses (Tobin, 2008; Tobin et al., 2008).

### The role of PKC in basal and U50,488H-induced KOPR phosphorylation

U50,488H-induced KOPR phosphorylation was inhibited to different degrees by GF109203X at all the four residues (Fig. 5A), by CHL at two residues (T363 and S369) (Fig. 5B), and by Go6976 at three residues (S356/T357 and S369) (Fig. 5C). GF109203X inhibits all PKC isoforms (conventional, novel and atypical PKCs) (Hofmann, 1997; Martiny-Baron et al., 1993), while CHL affects two PKC subfamilies (conventional and novel PKCs) (Herbert et al., 1990). Go6976 is a selective PKCα and PKCβ1 inhibitor. Thus, the three inhibitors reduced KOPR phosphorylation at the same residues to different degrees, which is likely due to their diverse effects on PKC isoforms and different extents of inhibition on each isoform. In addition, the findings that GF109203X, but not CHL and Go6976, reduced basal KOPR phosphorylation at S356/T357 indicate that atypical PKC isoforms play roles in basal S356/T357 phosphorylation. PKC has been reported to be involved in basal phosphorylation at S363 of MOPR and morphine-dependent MOPR phosphorylation (Chen et al., 2013; Hull et al., 2010; Johnson et al., 2006).

### PKC mediated agonist-independent KOPR phosphorylation

PMA, which activates conventional and novel PKCs, promoted KOPR phosphorylation at S356/T357, T363 and S369 (Fig. 6). PKC-mediated agonist-independent KOPR phosphorylation may be involved in heterologous desensitization of KOPR. PKC was reported to promote agonist-independent phosphorylation of other GPCRs, such as MOPR (Illing et al., 2014) and FFA4 receptor (Burns et al., 2014).

### Role of Gi/o proteins in U50,488H-induced KOPR phosphorylation

Pretreatment with PTX decreased U50,488H-stimulated KOPR phosphorylation at S356/T357, T363 and S369 (Fig. 2), indicating involvement of Gi/o proteins. The role of Gi/o proteins is likely related to their regulation of GRKs and PKC. Following G protein activation by GPCRs, GRKs 2 and 3 are recruited to plasma membranes by association with G_βγ_ proteins. PTX inhibits Gi/o_α_ proteins activation and thus blocks release of G_βγ_ subunits, resulting in no activation of GRKs 2 and 3. In addition, PKC activation is downstream of KOPR and G protein activation (Belcheva et al., 2005; Bohn et al., 2000). Thus, inactivation of Gi/o proteins may also impair PKC-mediated KOPR phosphorylation following U50,488H treatment.

### Effects of agonist-dependent and -independent KOPR phosphorylation on receptor internalization

Our results that GRKs2/3 and GRK6, but not PKC, are involved in U50,488H- induced KOPR internalization (Fig. 7) are consistent with previous reports. GRK2 and GRK3 are involved in U50,488H-induced internalization of the human and the rat KOPR (Li et al., 1999; McLaughlin et al., 2003; Schulz et al., 2002). The PKA/PKC inhibitor staurosporine did not inhibit U50,488H-induced rat KOPR internalization (McLaughlin et al., 2003).

We previously showed that U50,488H caused higher phosphorylation level at S356/T357, T363 and S369 and produced more double and triple phosphorylation, compared to etorphine. U50,488H induced KOPR internalization, but etorphine did not, indicating above-threshold KOPR phosphorylation is required for KOPR internalization (Chen et al., 2016). Knockdown of GRK2+3 and GRK6 affected all KOPR phosphoresidues more than PKC inhibitors (Fig.7). Thus, U50,488H-promoted PKC-mediated KOPR phosphorylation may be below the threshold necessary for internalization. PMA induced a low level of KOPR internalization, indicating PMA- promoted an above-threshold KOPR phosphorylation to trigger internalization.

Compared to U50,488H, PMA promoted much higher S356/T357 phosphorylation, much lower T363 phosphorylation and a similar level of S369 phosphorylation. In addition, PMA-promoted KOPR phosphorylation at S369 reached plateaus at later time than U50,488H (Fig. 5). Another possibility for lower internalization by PMA is phosphorylation rates. Thus, spatial and temporal regulations on KOPR phosphorylation may contribute to different cellular outcomes.

In summary, agonist-dependent (GRK- and PKC-mediated) and agonist-independent (PKC activation) KOPR phosphorylation yield different levels and rates of phosphorylation at each residue, leading to different internalization outputs.

### Role of receptor phosphorylation in KOPR signaling

Knockdown of GRKs increased U50,488H-induced ERK1/2 activation, suggesting that GRK-mediated KOPR phosphorylation desensitizes agonist-induced G protein-dependent ERK1/2 signaling. These results are similar to those of Doll et al (Doll et al., 2012) on the MOPR. In contrast, PKC-mediated KOPR phosphorylation is not involved in this desensitization. Taken together, GRK-mediated and PKC-mediated KOPR phosphorylation showed distinct regulations for desensitization of G protein-dependent signaling. One possible explanation is that GRK-, but not PKC-, mediated KOPR phosphorylation reaches a level to cause desensitization.

Our findings that GRKs and β-arrestins desensitized U50,488H-induced Gi/o-mediated ERK1/2 activation are consistent with those of Chavkin and colleagues that U69,593-induced rKOPR desensitization was regulated by GRK3, GRK5 and β-arrestin 2 (Appleyard et al., 1999).

Our result that PKC was not involved in U50,488H-induced ERK1/2 activation in N2A cells is different from the observations of Coscia and colleagues in C6 glioma and astrocytes (Belcheva et al., 2005; Bohn et al., 2000). The differences are likely due to cell systems used.

Our observation that U50,488H-induced ERK1/2 phosphorylation was G protein-dependent without β-arrestin involvement in N2A cells is similar to those of Jamshidi et al. (Jamshidi et al., 2015) in peripheral sensory neurons. In contrast, in astrocytes, activation of the KOPR by U69,593 yielded biphasic ERK1/2 activations: G protein-dependent activation in the early phase (at 5 min) and β-arrestin 2-dependent activation in the late phase (at 30 min and 2 hr) (McLennan et al., 2008). The differences may be due to different cell systems used.

## Acknowledgments

We thank Dr. Jeffrey Benovic and Dr. Walter Koch for providing reagents and Dr. Barrie Ashby for critical reading of the manuscript.

## Author contributions

Design experiments: Yi-Ting Chiu, Chongguang Chen, Stefan Schulz and Lee-Yuan Liu-Chen

Provide reagent: Stefan Schulz

Perform experiments: Yi-Ting Chiu and Chongguang Chen

Data Analysis: Yi-Ting Chiu, Chongguang Chen and Lee-Yuan Liu-Chen Write manuscript: Yi-Ting Chiu and Lee-Yuan Liu-Chen

## Supplementary data

**Table S1.**
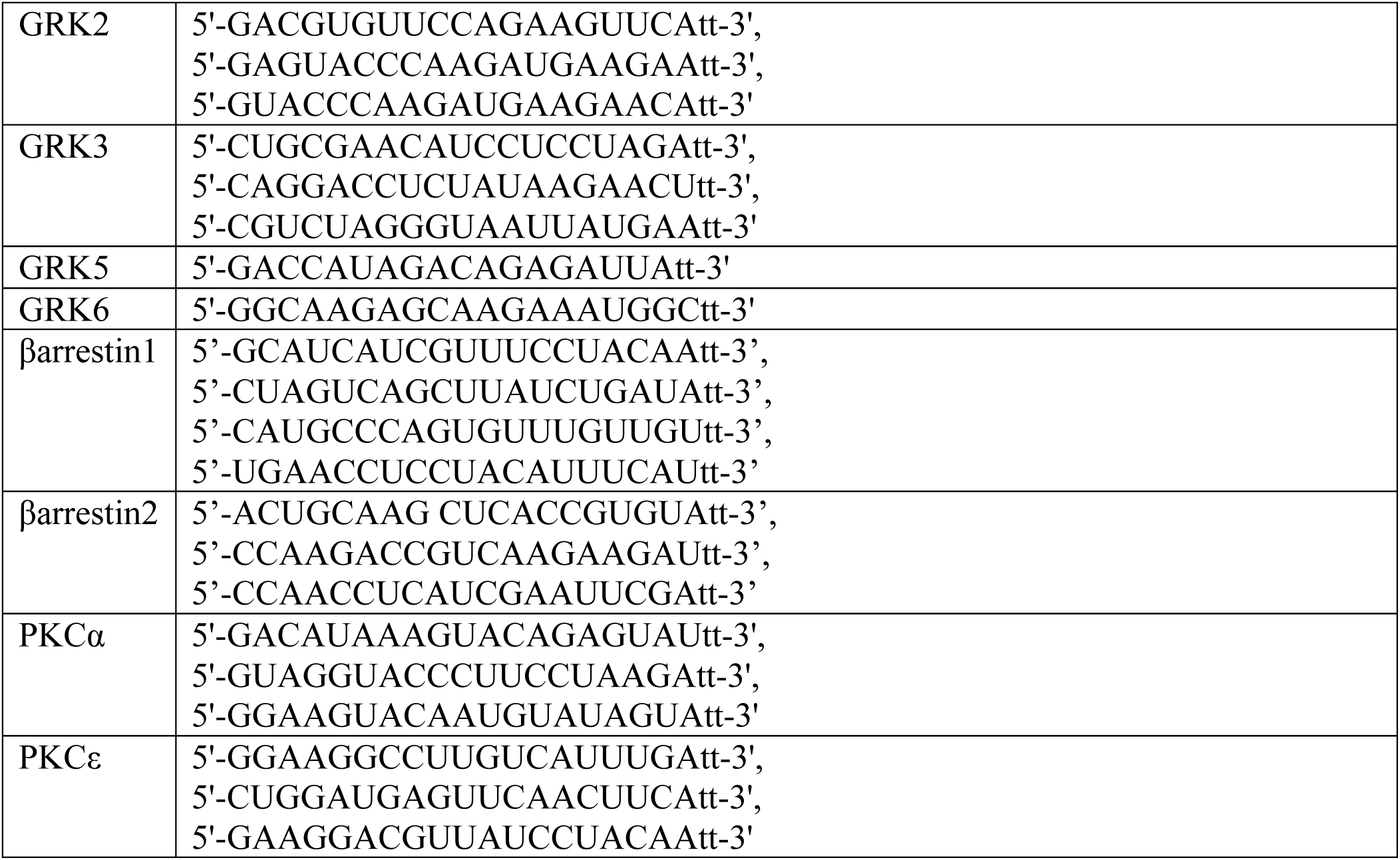
Sense sequences of all the siRNAs used in this study

**Figure S1.**
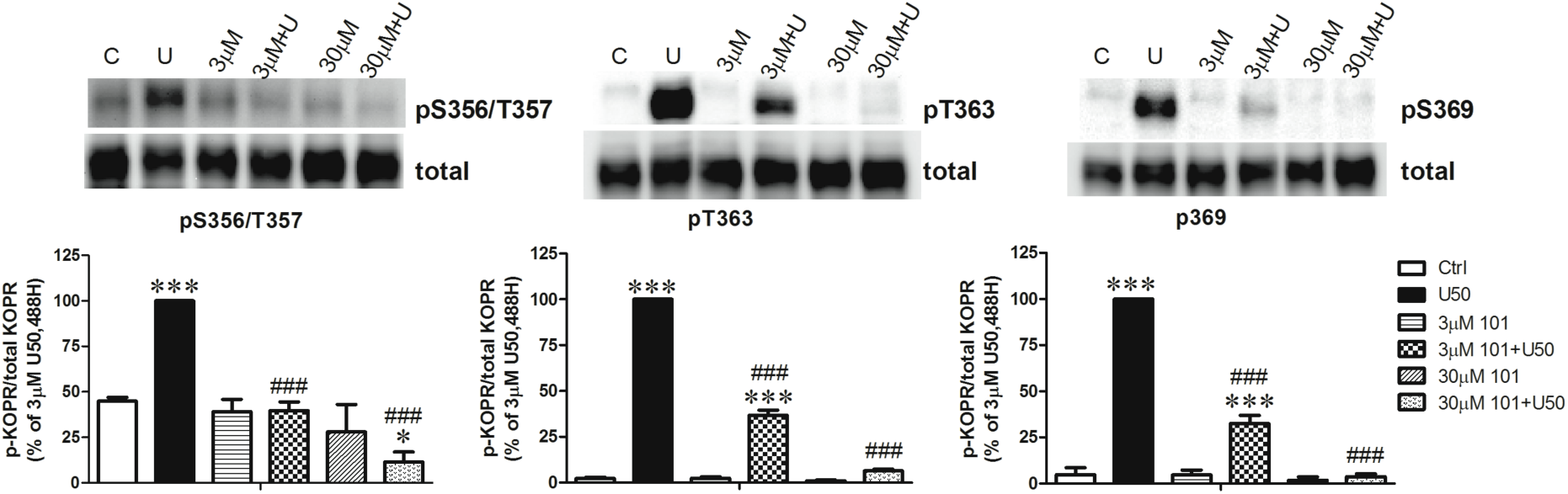
Effect of the GRK2/3 inhibitor compound101 on U50,488H-induced KOPR phosphorylation. Cells were pretreated with vehicle or compound 101 (3 or 30 µM) for 30 min followed by addition of 3 µM U50,488H for 2 min. KOPR phosphorylation was determined. The data shown are the mean ± S.E.M (n=3) and were analyzed by two-way ANOVA followed by Newman-Keuls *post-hoc* test (*: *p* <0.05, ***: *p* <0.001, compared to control group; ^###^: *p* <0.001, compared to U50,488H-treated group).

**Figure S2.**
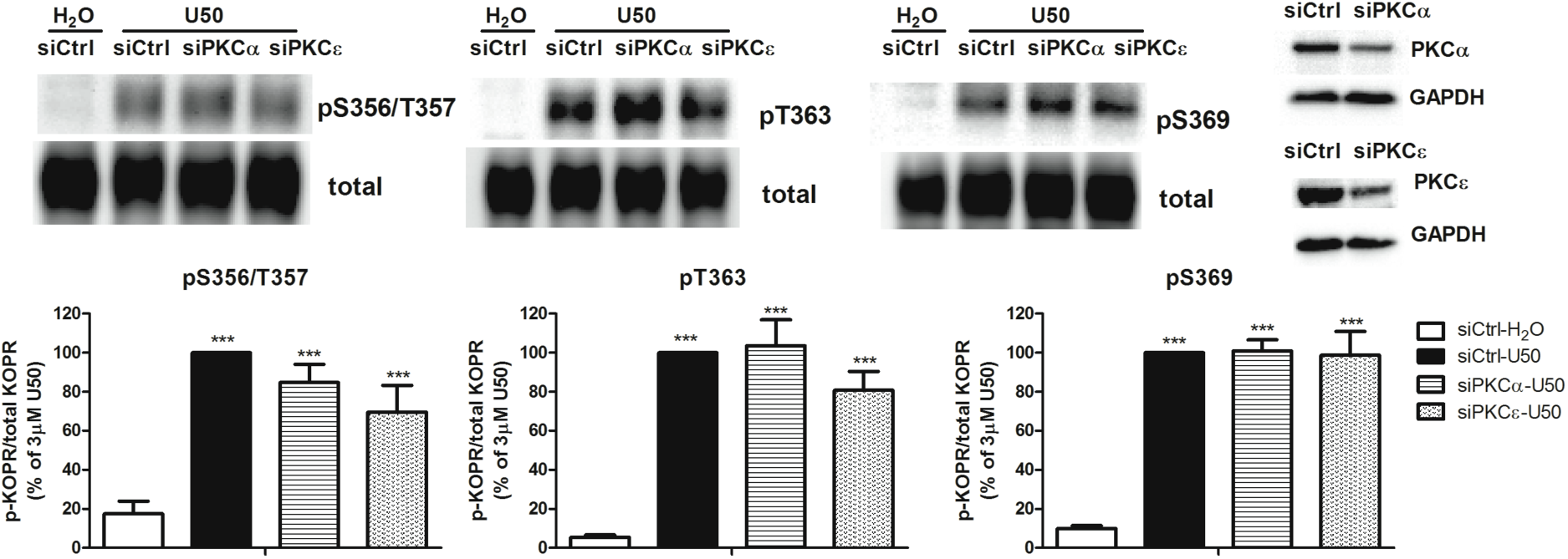
Effects of knockdown of PKCα and PKCε on U50,488H-induced KOPR phosphorylation. Cells were transfected with siRNAs targeting PKCα or PKCε (siPKCα and siPKCε, respectively) or the control siRNA (siCtrl) with Lipofectamine 2000. Two days later, cells were treated with vehicle or 3 µM U50,488H for 2 min and KOPR phosphorylation was determined. Each value presents the mean ± S.E.M (n=5). The data were analyzed by one-way ANOVA followed by Newman-Keuls *post-hoc* test (***: *p* <0.001, compared to vehicle control group). Knockdown of PKCα and PKCε were confirmed by immunoblotting.

